# A bioorthogonal chemistry approach to detect the K1 polysialic acid capsule in *Escherichia coli*

**DOI:** 10.1101/2022.11.04.515169

**Authors:** Vincent Rigolot, Yannick Rossez, Christophe Biot, Cédric Lion

## Abstract

Most Escherichia coli strains associated with neonatal meningitis express the K1 capsule, a sialic acid polysaccharide that is directly related to their pathogenicity. Metabolic oligosaccharide engineering (MOE) has mostly been developed in eukaryotes, but has also been successfully applied to the study of several oligosaccharides or polysaccharides constitutive of the bacterial cell wall. However, bacterial capsules are seldom targeted despite their important role as virulence factors, and the K1 polysialic acid (PSA) antigen that shields bacteria from the immune system still remains untackled. Herein, we report a fluorescence microplate assay that allows the fast and facile detection of K1 capsules with an approach that combines MOE and bioorthogonal chemistry. We exploit the incorporation of synthetic analogues of N-acetylmannosamine or N-acetylneuraminic acid, metabolic precursors of PSA, and copper-catalysed azide-alkyne cycloaddition (CuAAC) as the click chemistry reaction to specifically label the modified K1 antigen with a fluorophore. The method was optimized, validated by capsule purification and fluorescence microscopy, and applied to the detection of whole encapsulated bacteria in a miniaturized assay. We observe that analogues of ManNAc are readily incorporated into the capsule while those of Neu5Ac are less efficiently metabolized, which provides useful information regarding the capsule biosynthetic pathways and the promiscuity of the enzymes involved. Moreover, this microplate assay is transferable to screening approaches and may provide a platform to identify novel capsule-targeted antibiotics that would circumvent resistance issues.

## Introduction

Although *Escherichia coli* is an important part of the commensal microbiota colonizing the digestive tract of humans and animals, certain strains of this species have developed virulence attributes and cause serious diseases. Among these pathogenic strains, the meningitis/sepsis-associated *E. coli* (MNEC) pathotype is an important cause of neonatal infections. Mostly transmitted from mother to infant during birth, they are known to migrate to the vascular system after mucosal colonization, then penetrate the blood-brain barrier. This generally leads to meningitis or to septicaemia (blood-poisoning), both life-threatening conditions with high mortality and morbidity rates^1^. 80% of MNEC strains express the K1 capsule, a mucous layer of poly-*α*-2,8-sialic acid (PSA) that surrounds the bacterium thereby shielding its immunogenic proteins from detection by the host^2,3^. Mimicking the human glycan structure found in the neural cell adhesion molecule (NCAM), PSA is indeed not recognized as an external threat and allows the encapsulated bacteria to evade the immune system, and the severity of these infections is directly related to the amount of K1 antigen found at the bacterial surface^4–7^. This capsular polysaccharide is a linear homopolymeric chain constituted of *N*-acetylneuraminic acid (Neu5Ac) monomers belonging to the sialic acid (Sia) family of carbohydrates^8^. Its *de novo* biosynthesis requires the condensation of *N*-acetylmannosamine (ManNAc) and phosphoenolpyruvate to form Neu5Ac, which is then converted to the sugar nucleotide donor cytidine 5′-monophospho *N*-acetylneuraminic acid (CMP-Neu5Ac) and assembled into PSA in the cytosol prior to translocation to the cell surface^9^ **(Fig. 1)**. *E. coli* serotype K1, as well as other bacteria that produce polysialic acid capsules such as *Neisseria meningitidis, Pasteurella haemolytica* A2 or *Moraxella nonliquefaciens*, can also scavenge free Neu5Ac from the host^10^. Monosaccharide analogues modified with an additional chemical moiety can be used as molecular tools to engineer sialylated glycoconjugates in metabolic oligosaccharide engineering (MOE) approaches. Reutter and co-workers first pioneered MOE, with synthetic ManNAc analogues bearing an elongated *N*-acyl side chain that were successfully metabolized into cell surface sialoglycoproteins^11^. Modern labelling methodologies combine MOE and click chemistry or bioorthogonal chemistry, an ensemble of reactions developed by the groups of Sharpless, Meldal and Bertozzi, who were recently awarded the Nobel prize for this achievement^12–14^. A sugar reporter capable of entering the metabolic pathway under scrutiny and equipped with a reactive handle is first introduced in a living organism of interest, followed by the bioorthogonal ligation of an appropriately functionalized probe, most often a fluorescent dye for optical bioimaging^15^. These methods rely on biocompatible click reactions that involve pericyclic π_4_s + π_2_s processes, namely the copper-catalysed azide-alkyne (3+2) cycloaddition, the strain-promoted azide-alkyne (3+2) cycloaddition and the inverse electronic demand Diels-Alder reaction (CuAAC, SPAAC and IEDDA, respectively). Indeed, such mechanisms are mostly absent from living systems, thus greatly minimizing interference with biological processes^16^.

**Figure 1.**
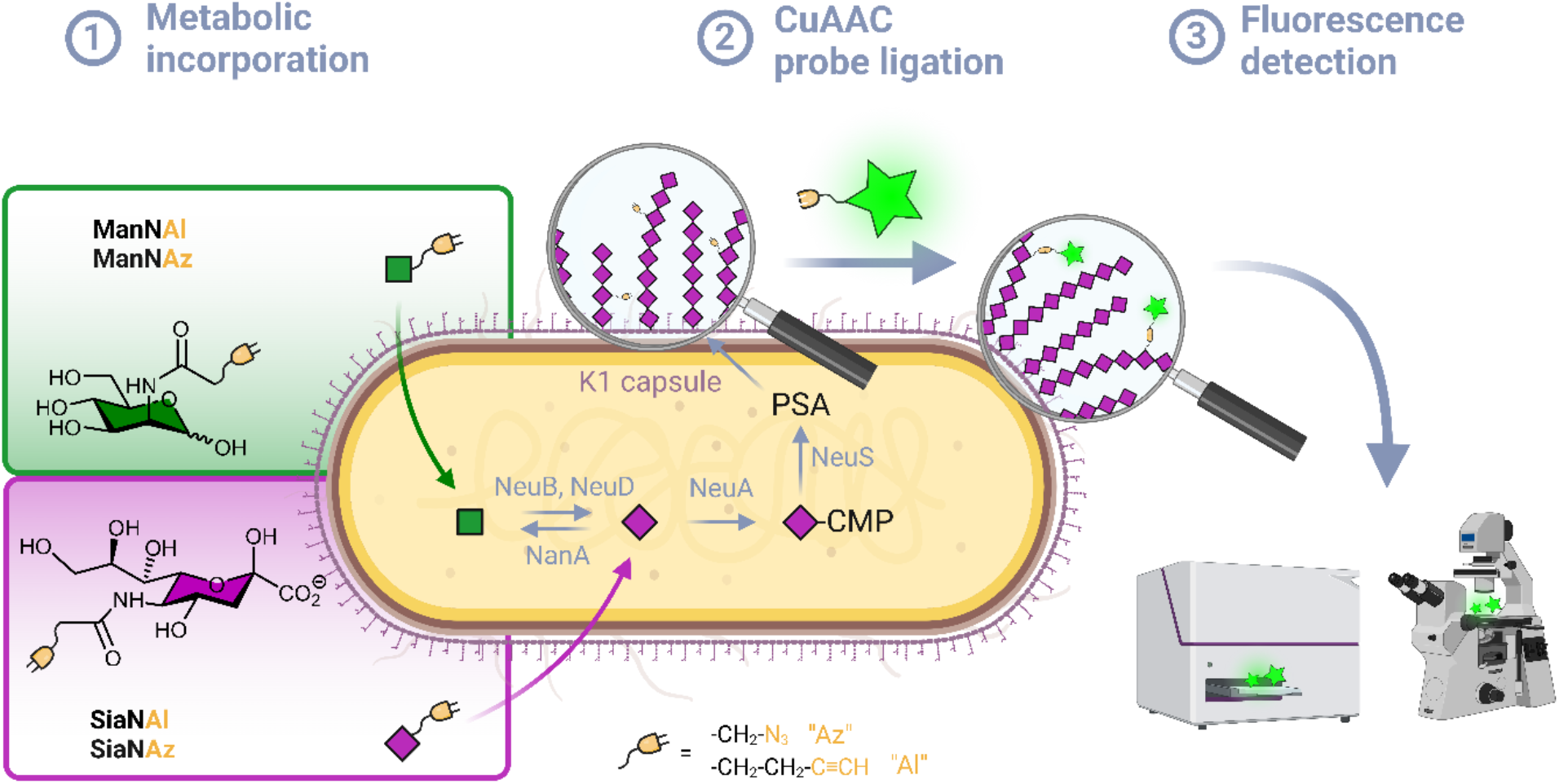
Labelling strategy workflow. **(1)** Chemical analogues of either ManNAc or Neu5Ac bearing an azide or alkyne moiety are incorporated in the capsule of living *E. coli* EV36 by hi-jacking the polysialic acid metabolic pathway. **(2)** The incorporated analogues are bioconjugated to a fluorescent probe *in vivo* by a CuAAC reaction. **(3)** The labelled bacteria are analysed with the desired technique such as fluorescence microscopy or fluorescence plate reader.

Although many bioorthogonal methods have been developed in prokaryotic cells to track proteins with non-canonical amino acids^17^, efforts to apply MOE to detect glycans in prokaryotes remain rather meager by comparison, as these strategies were developed in human models first and foremost. Some glycoengineering methods were reported in various species to label bacterial glycans, which were extensively reviewed by Banahene *et al*.^18^ They aim at detecting components of bacterial cell walls, such as peptidoglycans with *N*-acetylmuramic acid derivatives^19,20^ or lipopolysaccharides with, for example, azide-modified analogues of *N*-acetylglucosamine^21^, 3-deoxy-D-manno-octulosonic acid^22^ or legionaminic acid precursors^23^. However, very few bacterial species have been specifically labelled on their capsular polysaccharides by bioorthogonal chemistry. We could only identify two reports, in which capsules of commensal bacteria of the *Bacteroides* genus have been effectively tagged by MOE with modified *N*-acetylgalactosamine reporters^24,25^. The authors visualized the capsular component of prelabelled bacteria in live mice intestines, opening an avenue to better understand host-pathogen interactions and the evolution of related diseases.

To the best of our knowledge, capsular polysaccharides constituted of sialic acids have thus never been investigated by MOE in combination with click chemistry, despite the importance of these structures as virulence factors. Herein, we report a bioorthogonal labelling method to detect the polysialic acid capsule using alkyne- and azide-modified ManNAc and Neu5Ac reporters in *E. coli* K1 strains. Although this reporter toolbox is commonly used in mammalian systems to label *N*-glycoproteins^26,27^, it is the first time that these reporters are used successfully in *E. coli*. We aimed to take advantage of the fact that, unlike non-pathogenic strains, K1 strains are capable of synthesizing Neu5Ac *de novo*, and developed a test that allows facile and specific detection of the PSA capsule. The method was miniaturized into a microplate assay for transferability to screening approaches, which could serve as an interesting platform to deciphering K1 pathways and identifying external factors that influence PSA biosynthesis. Such assays could help in the discovery of new capsule-targeted classes of antibiotic drugs that decrease the virulence of a pathogenic bacterium while circumventing known resistance mechanisms.

## Results and discussion

### Chemical reporter

Per-*O*-acetylated monosaccharides are commonly used for MOE in mammalian models. However, per-*O*-acetylation can cause issues as it may generate non-specific signal and false positives^28,29^. It may also be conducive to acidification of the intracellular milieu, metabolic perturbation due to partially acetylated forms, or cytotoxicity.^16^ In addition, it has been suggested that low levels of non-specific esterase activity prevents the efficient release of the free sugar form and thwarts the use of such reporters in some prokaryotic species including *E. coli*^23^. Conversely, this species is capable of active transport mediated ManNAc uptake via the ManXYZ transporter and can also scavenge exogenous sialic acid from the host via the sialic acid transporter NanT. We therefore decided to use unprotected reporters and synthesized *N*-(2-azidoacetyl)-D-mannosamine (ManNAz), *N*-4-pentynoyl-D-mannosamine (ManNAl), *N*-(2-azidoacetyl)-neuraminic acid (SiaNAz) and *N*-4-pentynoylneuraminic acid (SiaNAl) (see experimental section). CuAAC was chosen as the bioorthogonal reaction, owing to its fast kinetics and ease of use. In addition, the azide and terminal alkyne tags are easily interchanged thus allowing comparison of the chemical handle’s impact.

### CuAAC whole cell labelling of K1 expressing E. coli EV36

We first assessed whether these reporters could be taken up and metabolically incorporated into living *E. coli* EV36, a strain containing all fourteen genes of the *kps* PSA biosynthetic cluster that produces the K1 capsule^30–32^. EV36 cultures were incubated overnight in lysogeny broth (LB) medium complemented with the modified monosaccharides, prior to CuAAC labelling with an appropriate clickable tetramethylrhodamine (TAMRA) dye bearing an alkyne or azide reactive moiety. The assay was implemented using a fluorescence microplate reader to enable facile testing of multiple conditions. The bioorthogonal ligation was first optimized to obtain a robust labelling and satisfying signal-to-noise ratio in EV36 whole-cells. Various parameters were screened, with TAMRA dye concentrations ranging from 625nM to 25μM in EV36 cultures incubated with reporters at a fixed 1mM concentration **(Fig S1)**, giving the best signal to noise ratio at 0.25μM of fluorophore for 45 minutes **(Fig 2A)**, and then with chemical reporter concentrations ranging from 100μM to 2mM at a fixed 0.25μM TAMRA concentration, resulting in an optimal chemical reporter concentration found at 600 μM **(Fig 2B)**.

**Fig 2.**
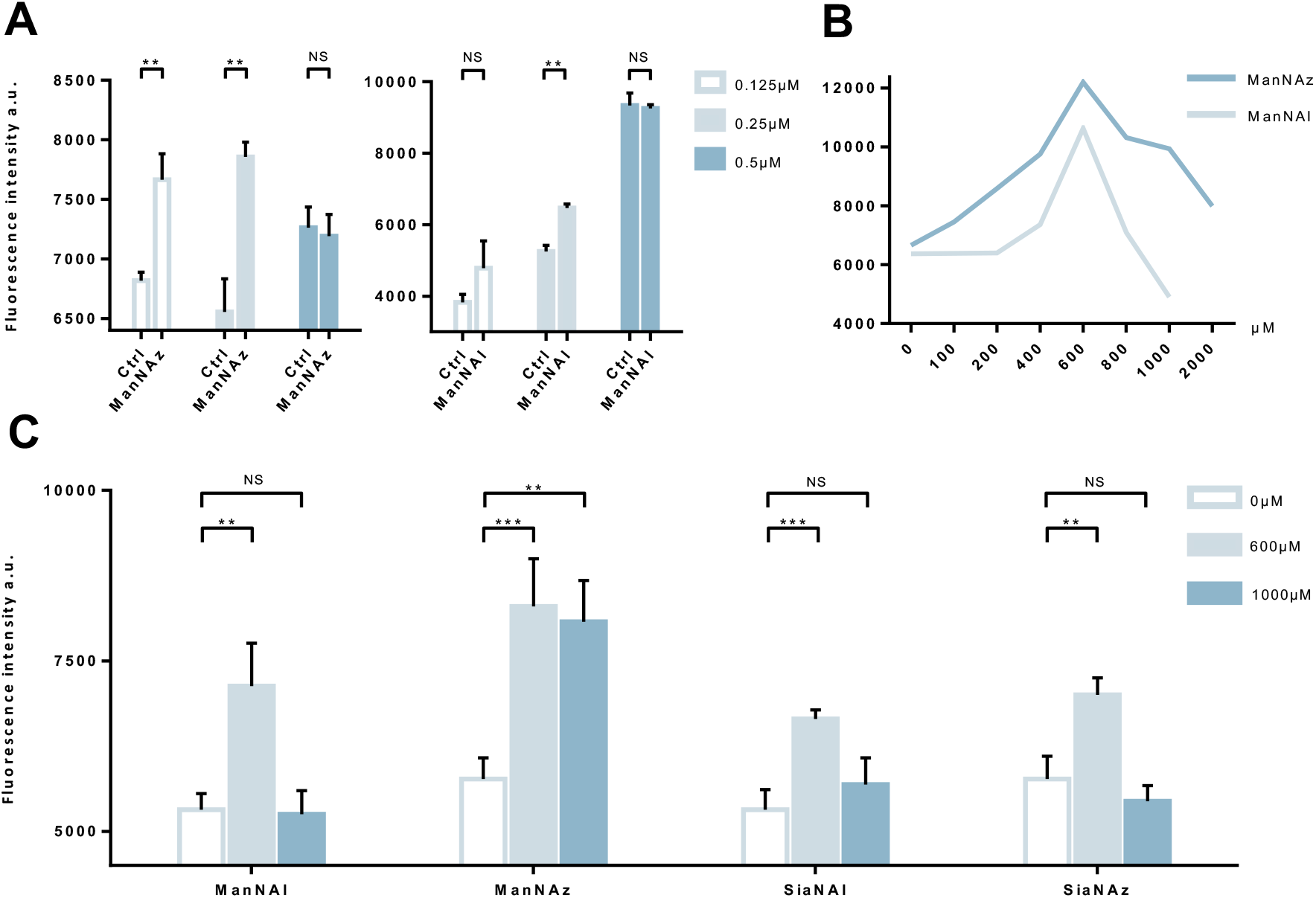
Whole cell suspension fluorescence after MOE in *E. coli* EV36. **(A)** Optimization of the fluorophore concentration. *E. coli* EV36 were grown with 1mM ManNAz, ManNAl or ManNAc (Control) followed by CuAAC ligation of TAMRA-Alk or TAMRA-N_3_ at 0.125, 0.25μM or 0.5μM. **(B)** Optimization of the chemical reporter concentration. *E. coli* EV36 were grown with ManNAl or ManNAz at concentrations ranging from 0 to 2000μM. CuAAC reaction was then performed with the fluorophore at 0.25μM. **(C)** Miniaturized assay in 96-well plates. 3 biological replicates with a minimum of 3 technical replicates were measured. Results are presented as mean + SD. Unpaired t-tests were performed to get statistical significance of difference observed compared to control condition (NS = non significant; ** = P < 0.01; *** = P < 0.001).

Next, we sought to perform statistically robust tests to determine the significance of the difference in labelling after the incorporation of the chemical reporters at 600μM and 1mM. In order to allow transferability of the protocol to prospective screening approaches, we developed this assay in 96-well plates, using an intensity-based readout with a fluorescence microplate reader. For all four compounds, we observe significant labelling at 600μM but not at 1mM, except for ManNAz being the only analogue where the labelling was significant in both cases **(Fig. 2C)**.

Experiments in which the bacteria were labelled with SiaNAz and SiaNAl generally exhibited a lower fluorescence intensity when compared to ManNAz and ManNAl, which came as a surprise. Indeed, these microbial cells are generally thought to be able to scavenge, process and install Sia derivatives but to metabolize ManNAc very slowly^33,34^, although this is highly species dependent. ManNAz led to the most efficient labelling. The noticeable difference in intensity between ManNAz and ManNAl tagging indicates that the azide handle may be more compatible with the metabolic network overall, or that the alkyne moiety might induce toxicity during the metabolic incorporation step. Distinct uptake and/or metabolisms should account for differences in labelling efficiency between ManNAc or Sia derivatives bearing the same chemical tag (*e*.*g*., ManNAz and SiaNAz)^33^.

### Reporter accumulation in the capsule

We pursued by investigating whether the observed labelling originates from reporters incorporated into the K1 capsule. To show this, *E. coli* EV36 was grown in LB medium complemented with ManNAz or ManNAc, then the K1 capsular polysaccharides were extracted and purified according to a previously described procedure^35^. A polyacrylamide gel electrophoresis of the extracted K1 capsule was performed, followed by alcian blue staining (a dye that specifically reveals acidic polysaccharides^36^) to confirm the presence and purity of PSA **(Fig. 3A)**. To evaluate the incorporation of the chemical reporters in the isolated capsule, the extracted acidic polysaccharides were then labelled in solution by CuAAC and fluorescence intensity was measured as described hereabove. Specific labelling was observed with both ManNAc derivatives ManNAl and ManNAz, but not with the Neu5Ac analogues SiaNAl and SiaNAz **(Fig. 3B)**. Therefore, this indicates that chemical reporters ManNAz and ManNAl enter the bacterial cell (possibly through the ManXYZ-encoded transporter) and are converted to SiaNAz and SiaNAl, respectively, in the bacterial cytoplasmic compartment^37^. The derived tagged Sia is subsequently activated as CMP-Sia, polymerized and exported at the cell surface by various enzymes and transporters^9^. *E. coli* has been described to convert ManNAc to ManNAc-6-P after ManNAc uptake to redirect the product on the degradation pathway^37,38^. Our results suggest that *E. coli* can also incorporate ManNAc into the K1 capsule biosynthesis *via* an unidentified pathway able to either transport ManNAc via the manXYZ phosphotransferase system, producing intracellular ManNAc-6-P which is then converted to ManNAc by an unknown phosphatase, or to transport ManNAc without ManNAc-6-P conversion. The former hypothesis carries a much stronger weight as the presence of this enzyme was previously suggested in *Neisseria meningitidis*, which also expresses K1 capsules, though no proper characterization has yet been reported^39^. Furthermore, Neu5Ac is transported by the NanT sialate permease to the interior of the bacterial cells and is known to be directly degraded^30,33,38,40,41^. Our results are in good agreement with this observation. Neu5Ac is indeed transported inside the cell as demonstrated by the whole cell fluorescence observed **(Fig. 2)**, but the monosaccharide is not directed to the PSA biosynthesis as suggested by the lack of signal detected on the purified capsule **(Fig. 3B)**.

**Fig 3.**
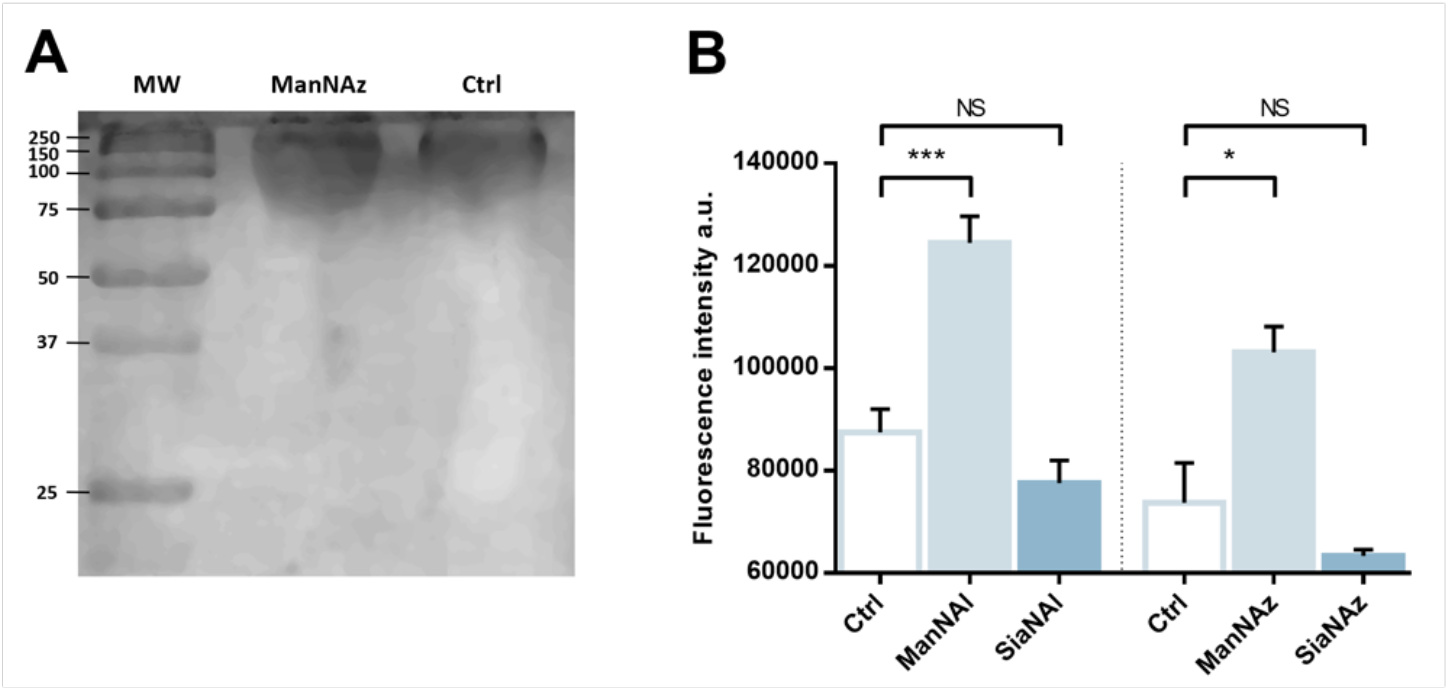
Capsular extraction and fluorescence quantification – **(A)** The capsular polysaccharides were purified after metabolic incorporation and analysed by PAGE with alcian blue staining. Molecular weight (MW) standard is indicated in kDa. **(B)** Capsular extracts obtained after metabolic incorporation of chemical reporters were labelled by CuAAC with 5μM TAMRA-N_3_ or TAMRA-Alk. 3 biological replicates with a minimum of 3 technical replicates were measured. Results are presented as mean + SD. Unpaired *t*-tests were performed to get statistical significance of difference observed compared to control condition (NS = non significant; * = P < 0.05; *** = P < 0.001).

To check whether constitutive biomolecules other than the capsule were labelled by our strategy, we carried out the same experiments on whole *E. coli* BL21 cells, a B strain that does not express capsules. All four reporters were tested, and no labelling was observed on this strain **(Fig. S2)**, thereby confirming the specific incorporation of the reporters into the K1 PSA capsule in the EV36 bacterium.

### Effect of unnatural monosaccharides on the K1 capsule biosynthesis

To ensure that the capsule was correctly expressed in the different labelling conditions, we performed immunofluorescence labelling of whole *E. coli* EV36 cells with an anti-K1 antibody followed by observation in fluorescence microscopy.

A neat peripheral labelling pattern was observed, confirming the presence and integrity of capsular PSA. The experiments were carried out after growth in the presence of a ManNAc or Neu5Ac analogue (ManNAz and SiaNAl, respectively), or after the thorough rinsing and shaking conditions required during the CuAAC labelling protocol (“Shaken” condition) **(Fig. 4)**. These results demonstrate a robust expression of the K1 capsule, without side effects of the unnatural monosaccharides on the K1 pathways.

**Figure 4.**
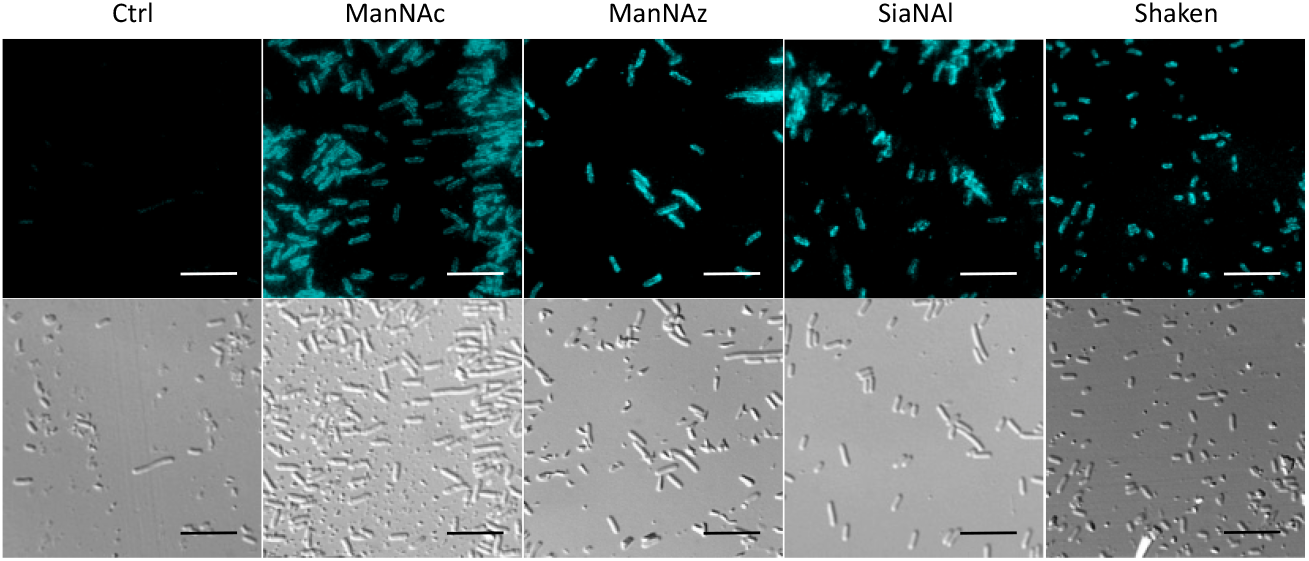
Immunofluorescence microscopy of the K1 capsule from *E. coli* EV36 with antibodies. *E. coli* EV36 was grown with either ManNAc, ManNAz, or SiaNAl 1mM before being incubated with an antibody directed against the K1 capsule followed by a fluorescent secondary antibody. Top row = fluorescence channel; bottom row= Differential interference contrast (DIC) channel; scale bar = 10μm. For the shaken condition, cells were submitted to the same shaking and rinsing conditions needed for CuAAC to confirm that the K1 capsule is still intact after these treatments. For the control condition, the secondary antibody was omitted to get the autofluorescence background.

### Impact of unnatural monosaccharides on the growth and physiological state of the bacteria

The *E. coli* EV36 growth and viability were evaluated for ManNAz, ManNAl, SiaNAz, or SiaNAl at 1 mM and 0.6 mM in comparison to ManNAc and LB alone. To evaluate the impact of the chemical reporters on the bacterial growth we measured the optical density at 600nm (OD_600_) of LB suspensions at regular intervals over the course of several hours. No difference was observed between the various conditions, illustrating that the reporters do not impair cell growth **(Fig. 5A)**. To evaluate the impact on long-term cell viability, serial dilutions of the suspensions were carried out after growth that were plated for viable counts. All chemical reporters except ManNAz induced rather significant toxicity on the long term when compared to ManNAc. Interestingly, when ManNAc was added to the medium the cell viability was also significantly lower by comparison to LB **(Fig. 5B)**. ManNAz and ManNAc equally affect bacterial viability to a mild extent. On the contrary, ManNAl drastically reduces the viability at both concentrations, revealing an increased toxicity linked to the appended pentynoyl moiety. Such an effect due to the nature of the bioorthogonal handle has been evidenced before in human Jurkat cells, where azidofucose derivatives proved to be more efficiently incorporated but more toxic than their alkynyl counterparts^42^. In the case of K1-expressing *E. coli*, it is the alkyne tag that decreases viability of EV36 cells, which might be attributed to a lower level of incorporation in capsular PSA leading to increased catabolism and accumulation of alkyne by-products interfering with other metabolic networks in the cell. Nevertheless, ManNAl is efficiently incorporated into the capsule **(Fig. 3B)**. It could thus be hypothesized that the pentynoyl sidechain is less compatible with the *E. coli* NeuB-encoded sialic acid synthase or NeuA-encoded CMP-sialic acid synthase than its 2-azido counterpart in ManNAz. This result calls attention to the use of unnatural monosaccharides to study bacterial physiology. To avoid any misinterpretation, viability assays should always be performed when testing MOE strategies on a new bacterial strain.

**Figure 5.**
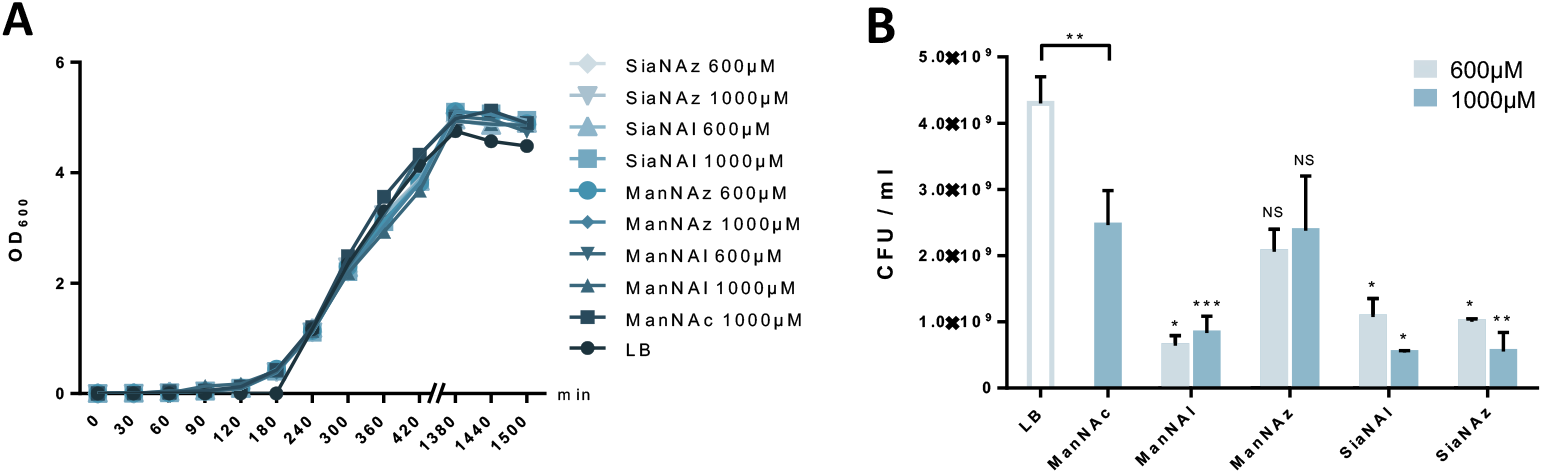
Growth and viability of *E. coli* EV36 when cultured with the unnatural monosaccharides. (**A**) Growth of *E. coli* EV36 cultures with different chemical reporters. (**B**) Viability assay of *E. coli* EV36 cultures after growth with different chemical reporters at 0.6mM and 1mM. 3 biological replicates with a minimum of 3 technical replicates were measured. Results are presented as mean + SD. Unpaired *t-*tests were performed to get statistical significance of difference observed compared to ManNAc 1mM condition (above error bars), as well as between ManNAc 1mM and LB (above brackets) (NS = non significant; * = P < 0.05; ** = P < 0.01; *** = P < 0.001).

### MOE microscopy

Given that ManNAz enables specific labelling of the capsule with no significant toxicity nor impact on growth, we chose to use this analogue for MOE CuAAC labelling of the capsule for observation by fluorescence microscopy. After ManNAz incorporation, cells were treated with a CuAAC buffer whose composition was adapted from a previously reported procedure^22^. A bright, heterogeneous and specific labelling was observed after ManNAz incorporation compared to ManNAc as negative control **(Fig 6)**. All cells appear to incorporate the reporter. In most cells, the labelling is of rather low intensity (although significant when compared to the control) but on some others, a bright peripheric fluorescent signal was visible **(Fig. 6B-C)**, typical of capsule localisation. Additionally, a number of cells exert a labelling pattern at the poles of the bacterium **(Fig. 6D)**, and most cells that do appear to be undergoing or to have just undergone cell division. This could be interpreted as a physiological state in which the older labelled PSA capsule has been relocated at the polar regions while new unlabelled capsule is being produced at the septum region during binary fission. Like most capsular polysaccharides in Gram-negative bacteria, PSA assembly and export through the periplasm is controlled by a multiprotein complex. The K1 antigen belongs to group 2, which identifies strains for which this export involves an ABC-transporter^43^. Our result might indicate that this translocation to the outer membrane is spatially regulated at the equatorial region during cell division, similarly to what has been proposed for Gram-positive *Streptococcus pneumoniae*, whose capsule export is mediated by the Wzx-Wzy system^44^. Indeed, Henrique *et al*. suggested that this spatial control ensures the coordination of capsule and cell wall biogenesis at the division site, leaving the bacteria unexposed to the host’s immune system throughout. The heterogeneous labelling points out the incorporation of ManNAz with a potential dependence on the physiological state of the bacteria during their growth. EV36 bacteria continue to grow and multiply post-labelling, resulting in signal dilution over time for bacteria that are dividing after CuAAC treatment. Similarly, the bacteria need to adjust their metabolism to ManNAc and ManNAz import and successfully export the polysialic acid at the cell surface, which may also result in heterogeneity. Finer localisation may be obtained with super-resolution microscopy, to refine the visualization of capsule export zones at the poles of the cells.^45^

**Fig 6.**
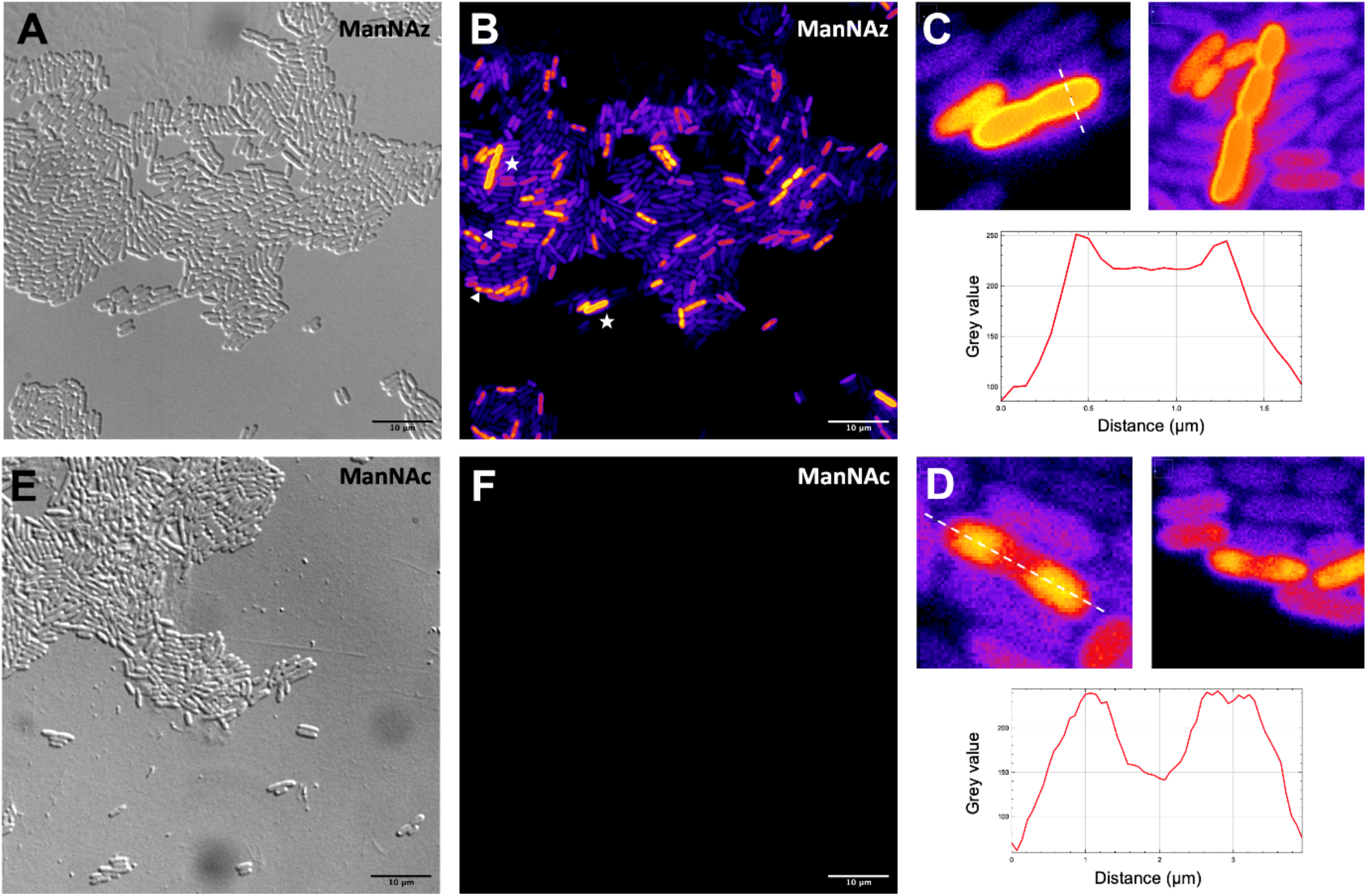
Fluorescence microscopy of *E. coli* EV36 after incorporation of ManNAz **– (A, B)** Incorporation of ManNAz in E. coli EV36, brightfield in gray, fluorescence channel in “Fire” colorscale. **(C)** (top) Magnified zooms on cells showing a pericellular labelling (highlighted by white stars in B insert) and (bottom) intensity profile along the transversal dashed line; **(D)** (top) Magnified zooms on cells showing a polar labelling (highlighted by white triangles in B insert) and (bottom) intensity profile along the longitudinal dashed line. (E, F) Negative control incorporating unmarked ManNAc. Experiments were carried out as 3 biological replicates. Scale bars= 10μm.

In order to further confirm the specificity of our method for the K1 capsule and ascertain its applicability to various pathogenic bacteria, we then tested three other wild-type *E. coli* strains. In addition to *E. coli* BL21, a strain devoid of capsule mentioned hereabove as negative control **(Fig S2)**, we also evaluated the ability of *E. coli* Nissle 1917 bacteria to incorporate ManNAz. The Nissle 1917 strain expresses another capsular acidic polysaccharide, namely the K5 antigen heparosan constituted of a disaccharide repeating unit that contains glucuronic acid and *N*-acetylglucosamine. This strain was unable to incorporate ManNAz, as depicted in **Fig. 7**, further confirming the specificity of this assay for the K1 PSA capsule. Moreover, the two *E. coli* K1 pathogenic strains K-235 and U5/41, which produce significantly higher levels of PSA than the EV36 model, exhibited strong fluorescence **(Fig7)**. This indicates an increased level of incorporation of SiaNAz units into PSA and derived from the metabolization of ManNAz when compared to experiments on EV36. The heterogeneity was lesser in the K-235 and U5/41 wild-types and the pattern observed was consistent, with bacteria marked at the periphery typical of capsule staining and other bacteria showing localization at the poles **(Fig S3)**. In contrast, the heterogenous incorporation observed in the EV36 construct might be attributed to a less efficient metabolic channeling process, induced by the expression of exogenous genes from K1 strains in a K-12 background (*e*.*g*., the *manXYZ* operon).

**Fig 7.**
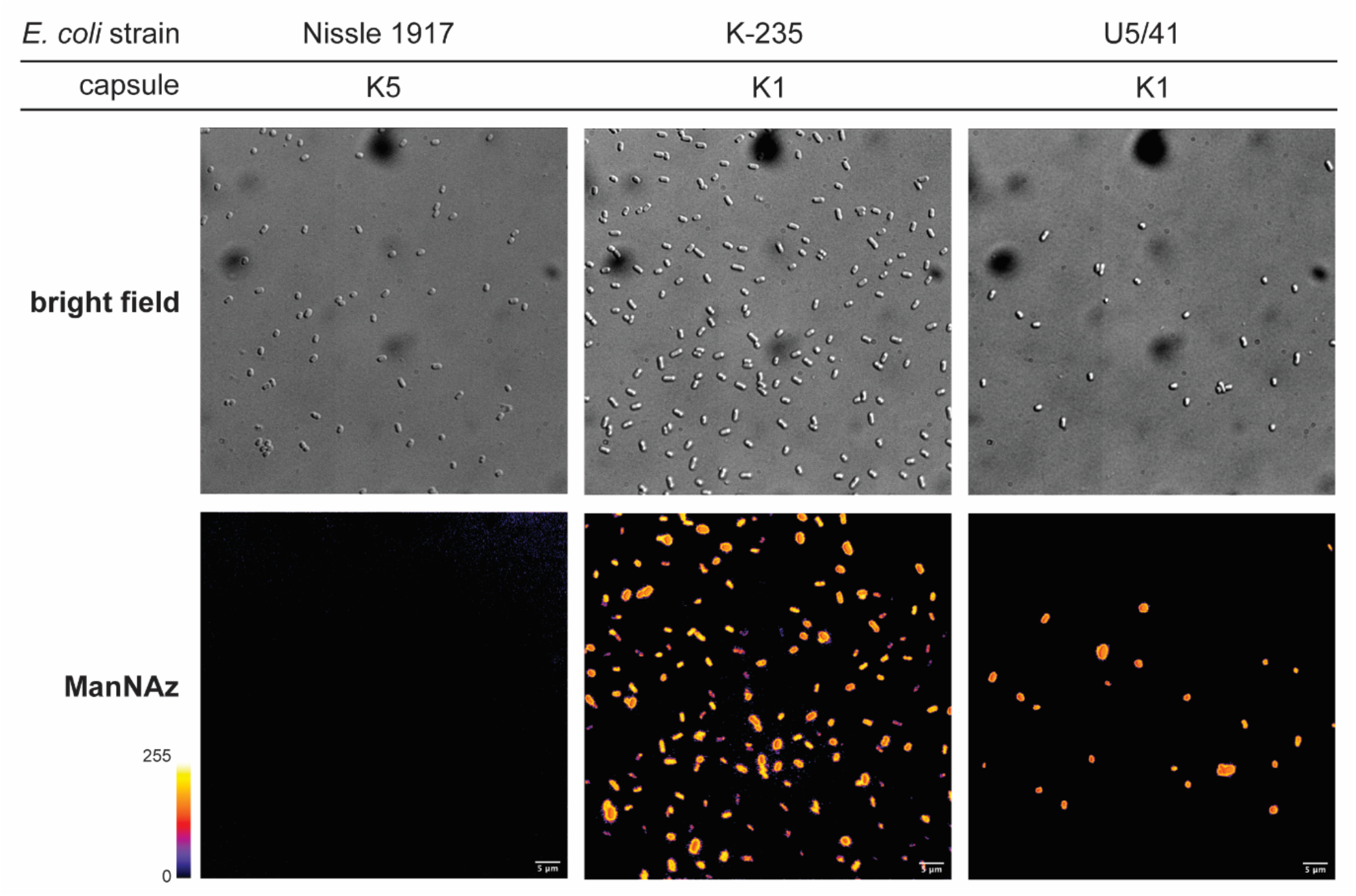
Fluorescence microscopy of *E. coli* Nissle 1917 expressing K5 capsule, and K-235 and U5/41 expressing K1 capsule grown with ManNAz **– (top)** Brightfield channel; **(bottom)** fluorescence channel in Fire colorscale. Experiments were carried out as 3 replicates. Scale bar = 5 μm.

Thus, our method can be used to detect K1 antigen production in pathogenic strains, which could help decipher the dynamics of capsule expression and the factors that regulate it in future studies. Furthermore, it provides a platform for screening new types of antibiotics targeting K1 capsule metabolism.

## Experimental procedures

### Chemical synthesis of unnatural monosaccharides

ManNAl (*N*-4-pentynoyl-D-mannosamine): 4-pent-ynoïc acid (502 mg, 5.1 mmol, 1 eq.) was solved into CH_2_Cl_2_ (25 mL). *N*-hydroxysuccinimide (674 mg, 5.8 mmol, 1.14 eq.) and EDC or 1-ethyl (3-dimethylaminopropyl)carbodiimide (1.96 g, 10.2 mmol, 2eq.) were slowly added. The mixture was stirred for 3h30 at room temperature. The end of reaction was confirmed by TLC (cyclohexane/AcOEt; 70 : 30). The solution was then washed four times with KHSO4 (aqueous solution 2.8%). The recombinated organic layers were dried over sodium sulfate, filtered, and then concentrated under reduced pressure to give the succinimidyl ester as a white solid (875.3 mg, 4.48 mmol, 88 %). The product was used in the next step without further purification. The coupling reaction with Mannosamine hydrochloride was conducted under nitrogen atmosphere. Hydrochloride D-Mannosamine (759 mg, 3.52 mmol, 1 eq.) and the succinimidyl ester from previous step (688 mg, 3.52 mmol, 1 eq.) were solved into DMF (25 mL). Triethylamine (1.4 mL, 10.77 mmol, 3 eq.) was slowly added. The mixture was stirred overnight. DMF was then removed under reduced pressure and the crude product was purified by flash chromatography (silica column, 40 g, 30 μm) with an elution by CH_2_Cl_2_ / EtOAc / MeOH ; 45 : 45 : 10). Solvents were then eliminated to yield ManNAl as a white solid (830 mg, 3.34 mmol, 95 %). Mixture of anomers (α / β ≈ 60% / 40%).

<u>α ^1^H NMR (300 MHz, D_2_O):</u> δ = 5.00 (d, J=1.4, 1H, H_1_), 4.23 (dd, J=4.7, 1.4, 1H, H_2_), 3.93 (dd, J=9.8, 4.7, 1H, H_3_), 3.80 – 3.62 (m, 3H, H_5 &_ H_6_), 3.50 (t, J=9.6, 1H, H_4_), 2.58 – 2.32 (m, 4H, H_8 &_ H_9_), 2.26 (t, J=2.3, 1H, H_11_). ^13^C NMR (75 MHz, D_2_O): δ = 175.26 (C7), 93.17 (C1), 83.43 (C10), 71.94 (C5), 70.14 (C11), 68.73 (C3), 66.72 (C4), 60.36 (C6), 53.15 (C2), 34.10 (C8), 14.40 (C9).

<u>β ^1^H NMR (300 MHz, D_2_O):</u> δ = 4.90 (d, J=1.6, 1H, H_1_), 4.35 (dd, J=4.4, 1.4, 1H, H_2_), 3.82 – 3.61 (m, 3H, H_3 &_ H_6_), 3.40 (t, J=9.8, 1H, H_4_), 3.29 (ddd, J=9.9, 4.8, 2.3, 1H, H_5_), 2.53 – 2.31 (m, 4H, H_7 &_ H_8_), 2.26 (t, J=2.3, 1H, H_11_). ^13^C NMR (75 MHz, D_2_O): δ = 175.98 (C7), 92.84 (C1), 83.85 (C10), 76.28 (C5), 71.96 (C3), 69.99 (C11), 66.45 (C4), 60.36 (C6), 53.98 (C2), 34.25 (C8), 14.26 (C9).

SiaNAl (N-4-pentynoylneuraminic acid): ManNAl, (60 mg, 0.23 mmol, 1 eq.), sodium pyruvate (46 mg, 0.345 mmol, 2 eq.) and Neuraminic5Ac aldolase (Sigma Aldrich EC 4.1.3.3) from E. Coli K12 (5 units dissolved in 0.1 mL) were added into 600 μL of phosphate buffer (KH_2_PO_4_, 10.3 mM, K_2_HPO_4_, 52.8 mM and MgCl_2_ 20 mM, pH – 7.5) into a reaction tube which was incubated at 37.5°C with moderate shaking (140rpm) for 18 hours. The reaction completion was monitored by TLC (propan-1-ol / 25% Ammonia / H_2_O ; 6 : 1 : 2.5). Reaction was quenched by the addition of 30 mL of water. The product was purified on an anion exchange resin (Bio-Rad AG1X2) activated with NH_4_HCO_3_ 0.1M. The sample was loaded on the column which was washed with H2O (5 CV). Elution was performed with aqueous NH_4_HCO_3_ (0.05M, 5 CV) and then NH_4_HCO_3_ (0.2M, 5 CV). Fractions were detected by TLC (same elution as for the reaction monitoring) combined and freeze dried. After being solved into H2O (1mL), sample was passed through a gel filtration column (P2) and then freeze dried again. SiaNAl was obtained as white powder (73 mg, 0.21 mmol, 91 %).

^1^H NMR (600 MHz, D_2_O): δ = 3.96 – 3.86 (m, 2H, H_4 &_ H_9_), 3.83 (t, J=10.1, 1H, H_5_), 3.72 (dd, J=11.9, 2.7, 1H, H_9_), 3.64 (ddd, J=9.2, 6.7, 2.7, 1H, H_8_), 3.51 (d, J=9.2, 1H, H_7_), 3.49 – 3.43 (m, 1H, H_9’_), 2.47 – 2.34 (m, 4H, H_11 &_ H_12_), 2.28 (d, J=2.0, 1H, H_13_), 2.10 (dd, J=13.0, 4.9, 1H, H_3eq_), 1.70 (dd, J=12.7, 11.7, 1H, H_3ax_). ^13^C NMR (75 MHz, D_2_O): δ 175.43 (C1), 172.91 (C10), 95.16 (C2), 83.54 (C13), 70.40 (C14), 70.37 (C8), 70.23 (C6), 68.29 (C7), 66.45 (C4), 63.21 (C9), 52.04 (C5), 38.79 (C3), 34.67 (C11), 14.53 (C12). m/z: calculated: [M]+ = 346.312 ; measured: 346.017

ManNAz (*N*-azidoacetyl-D-mannosamine): synthesis, 2-azidoacetic acid (117mg, 1.16mmol, 1eq) was dissolved along with DIC (*N,N’*-diisopropylcarbodiimide, 175mg, 1.392mmol, 1.2eq), HOBt (hydroxybenzotriazole, 195mg, 1.276mmol, 1.1eq) and DIPEA (*N,N*-diisopropylethylamine, 141mg, 1.392mmol, 1.2eq) in DMF (dimethylformamide, 20ml), D-mannosamine hydrochloride (250mg, 1.16mmol, 1eq) was added and the reaction was stirred at room temperature for 19h under argon. The total consumption of mannosamine was assessed by silica thin layer chromatography (TLC) (CH_2_Cl_2_/MeOH 9:1 v/v) before solvent removal under reduced pressure. The reaction crude was purified by silica flash column chromatography (50μm, 40g, dry load, CH_2_Cl_2_/MeOH 95:5 v/v), and fractions containing the product were gathered and concentrated under reduced pressure, affording the ManNAz as a white powder (175mg, 0.67mmol, 57%).

^1^H NMR (300 MHz, D_2_O): δ = 5.10 (d, J=1.0 Hz, 1H, H_1_ α), 5.01 (d, J=1.4 Hz, 1H, H_1_ β), 4.46 (dd, J=4.0 Hz, 1.4Hz, 1H, H_2_ β), 4.33 (dd, J=4.4 Hz, 1.0 Hz, 1H, H_2_ α), 4.10 – 3.98 (m, 5H, H_3_ α + H_8_ αβ), 3.88 – 3.71 (m, 5H, H_5_ α + H_3_ β + H_6_ αβ), 3.56 (t, J=9.5Hz, 1H, H_4_ α), 3.46 (t, J=9.8Hz, 1H, H_4_ β), 3.38 (ddd, J=9.8Hz, 4.7Hz, 2.1Hz, 1H, H_5_ β). ^13^C NMR (75 MHz, D_2_O): δ = 171.87 (s, C7 α), 170.97 (s, C7 β), 92.89 (s, C1 α), 92.83 (s, C1 β), 76.40 (s, C5 β), 72.03 (s, C3 β), 71.97 (s, C5 α), 68.82 (s, C3 α), 66.76 (s, C4 α), 66.52 (s, C4 β), 60.41 (s, C6 β), 60.39 (s, C6 α), 54.28 (s, C2 β), 53.37 (s, C2 α), 51.72 (s, C8 β), 51.65 (s, C8 α).

SiaNAz (*N*-azidoacetylneuraminic acid): ManNAz (42mg, 0.159mmol, 1eq) was dissolved in 600μl PBS (pH 7.6) with sodium pyruvate (32mg, 0.239mmol, 1.5eq) in a 5ml capped tube. 5 units of Neu5Ac aldolase (Sigma Aldrich, EC 4.1.3.3) were added and the reaction was stirred at 37°C for 16h. The formation of the product was checked by TLC (1-propanol, NH_3_, H_2_O 6:1:2,5) with resorcinol as a reagent. The reaction crude was purified by anion exchange chromatography (BioRad, AG1X8; H_2_O 5CV, NH_4_HCO_3_ 0.05M 5CV, NH_4_HCO_3_ 0.2M 5CV). Fractions containing the product were identified by TLC as previously then gathered and concentrated under reduced pressure. The white powder obtained was desalted using size exclusion chromatography (P2) with ultrapure water as eluent. Fractions containing the product were gathered, concentrated under reduced pressure then freeze-dried, affording the product as a white powder (18.5mg, 0.053mmol, 34%).

^1^H NMR (300 MHz, D_2_O): δ 4.09-3.88 (m, 5H, H_5_+H_4_+H_3_+H_2_), δ 3.78 (dd, J=11.2Hz, 2.2Hz, 1H, H_8cis_), δ 3.65 (ddd, J=8.5Hz, 4.1Hz, 1.2Hz, 1H, H_7_), δ 3.55 (dd, J=11.2Hz, 5.8Hz, 1H, H_8trans_), δ 3.45 (dd, J=9.2Hz, 0.9Hz, 1H, H_6_), δ 2.18 (dd, J=12.5 Hz, 4.8Hz, 1H, H_1_ eq), δ 1,75 (dd, J=12.4Hz, 11.8Hz, 1H, H_1_ ax). ^13^C NMR (75 MHz, D_2_O): δ 70.8 (CH, C5), δ 69.82 (CH, C7), δ 68.43 (CH, C6), δ 67.01 (CH, C2), δ 63.10(CH2, C8), δ 52.21(CH, C3), δ 51.9 (CH2, C4), δ 39.32 (CH2, C1)

### Bacterial strains and growth conditions

Five bacterial strains were used in this study. The non-pathogenic model *E. coli* EV36^30^ that expresses the K1 capsule was kindly provided by Pr Antonia P. Sagona. The K1 producing pathogenic strains *E. coli* K-235 (ATCC 13027) and *E. coli* U5/41 (ATCC 11775) were purchased from ATCC. *E. coli* BL21^46^, a well-described B strain, and *E. coli* Nissle 1917 expressing the K5 capsule were used as negative controls to demonstrate the specificity of the method for K1 capsule and were kindly provided by Dr Marie Titecat. The bacteria were streaked out from -80°C stocks on LB agar Petri dishes and cultured overnight at 37°C. Isolated colonies grown on these plates were used to inoculate liquid cultures for the rest of the experiments.

### Metabolic incorporation and CuAAC buffer preparation

*E. coli* liquid cultures prepared as explained above were diluted to an OD_600_ of 1. 20μl of this solution was transferred to 10ml LB (supplemented with the desired chemical reporters) in a 50ml conical tube and grown overnight at 37°C/180rpm. For fluorescence labelling by CuAAC, fresh aqueous stock solutions of the different reagents were used to prepare the CuAAC buffer. It should be prepared several minutes to an hour prior to use and should not be kept for more than a few hours. CuSO_4_ (1mg/ml, final concentration 150μM) and BTTAA (10mg/ml, final concentration 300μM) are first mixed and added to the fluorescent probe (TAMRA-Alk or TAMRA-N_3_, 1mg/ml, desired final concentration). K_2_HPO_4_ (100mg/ml, final concentration 100mM) and H_2_O are then added. Finally, sodium ascorbate (10mg/ml, final concentration 2,5mM) is added right before use.

### Microplate fluorescence

Overnight cultures were adjusted to OD_600_ 1 by diluting the suspensions with PBS. 200μl of these suspensions were split in 2ml microtubes (minimum of 3 per condition) then centrifuged (2min, 10 000G), and pellets were resuspended in 200μl CuAAC buffer and agitated for 45min, 600rpm at room temperature in the dark. These suspensions were rinsed 3 times with 1ml PBS (2min, 10 000rpm) before being resuspended in 200μl PBS. CuAAC whole cell suspensions were split in a dark opaque 96-well plate (100μl/well). Fluorescence (λ_em_ 535±20nm/λ_exc_ 585±30nm) was measured on a CLARIOstar Plus microplate reader.

### Fluorescence microscopy

Agar pads were made by pouring 10μl of hot LB agar on a microscopy slide and quickly covering it with a glass coverslip. Whole-cell suspensions were deposited on the hardened agar pad by gently raising the coverslip and placing it back down. These slides were then observed on a Leica AF6000 LX inverted video microscope with differential interference contrast (DIC). For immunofluorescence staining of the capsule, overnight cultures were adjusted to 1 OD_600nm_ by diluting the suspensions with PBS and treated with anti-K1 rabbit antibody (ENZ-ABS559-0100 Enzo Life Sciences) (1/100, 45min), rinsed with 200μl PBS and treated with anti-rabbit 488 antibody (1/250, 30min), rinsed with 200μl PBS and finally mounted on agar pads and observed as described above. For CuAAC fluorescence labelling, cells were treated as described for the microplate fluorescence with the fluorophore concentration adjusted to 100mM. After the reaction and rinsing steps, cells were resuspended in 200μl PBS and 10μl of these suspensions were mounted on agar pad before observation on a fluorescence videomicroscope.

### Viability assay and growth curves

For viability assay, OD_600_ of overnight cultures was harmonized to 1 by diluting the suspensions with sterile PBS. 200μl of these suspensions were transferred in a 96-well plate. Serial dilutions ranging from 10^−1^ to 10^−10^ were performed. 20μl of these suspensions were streaked on LB agar Petri dishes and grown overnight at 37°C. Petri dishes showing between 5 to 250 colonies were selected, colonies counted, and the number of CFU in the original suspension was deducted from the dilution. For growth curves, from an isolated colony, 2ml of LB liquid culture were grown in a 15ml conical tube for 2 hours at 37°C. 10μl of this suspension was used to inoculate 10ml LB supplemented with the desired chemical reporter or monosaccharide in a 50ml conical tube. OD_600_ of these suspensions was then recorded at regular intervals.

### Capsule extraction and quantification

Capsular extracts were purified by following a method described previously^35^. Overnight grown cultures were suspended in 600μl lysis buffer (100mM SDS, 50mM Tris, 0.128mM NaCl). 600μl phenol/chloroform/isoamyl alcohol (25:24:1) were added and agitated 15min at 65°C. After centrifugation (16 000g for 15min at 4°C), the upper phase was transferred and completed with 600μl of ice-cold absolute ethanol. The tubes were stored overnight at -20°C. Tubes were then centrifuged (10min, 13000g) and the white precipitate was rinsed with ethanol then dried under N_2_ flow. 20μl DNAse was added (45min, 37°C, 300rpm) followed by 20μl Proteinase K (1h, 56°C, 300rpm). 560μl H_2_O and 600μl phenol/chloroform/isoamyl alcohol (25:24:1) were added. Tubes were centrifuged (15min, 13000rpm) and the upper phase transferred in a new microtube. 200μl H_2_O, 50μl sodium acetate 3M and 1m absolute ethanol were added and the tubes were stored at -20°C overnight. The tubes were centrifuged (15min, 13000rpm) and the white precipitate was rinsed with 1ml ice cold absolute ethanol and dried under N_2_ flow. Capsular extracts were finally resuspended in 50μl of H_2_O. The capsule extracted was labelled and quantified on a microplate reader as follow: 10μl of the capsular extract solutions were added to 190μl CuAAC buffer in a microtube and agitated 45min, 600G at RT. After the reaction 50μl sodium acetate 3M and 1ml ice cold absolute ethanol were added and tubes were stored overnight at -20°C. Tubes were then centrifuged (15min, 13 000G and the white pellet was rinsed with ice cold absolute ethanol, dried under N_2_ flow, resuspended in 20μl H_2_O and transferred to a dark opaque 96-well plate for fluorescence readout on a microplate reader. The purity of the capsular extract was performed by polyacrylamide gel electrophoresis, for this 10μl of the same capsular preparation were loaded on a 15% TBE-PAGE (Tris-Boric-EDTA polyacrylamide gel electrophoresis) as described previously^47^. The capsular polysaccharides were stained with 5% alcian blue in water for 15min followed by three washing steps with acetic acid 5% in a solution of 50% methanol until the gel background get unstained.

## Conclusions

We developed a method for the specific bioorthogonal labelling of K1 capsules in *E. coli* after the metabolic incorporation of ManNAc analogues equipped with alkyne and azide chemical handles. These sialylation reporters did not exert any significant effect on bacterial growth. ManNAz was determined as the better choice among the tested reporters, as ManNAl showed inherent long-term cytotoxicity. While both ManNAc and Neu5Ac derivatives were readily incorporated in growing *E. coli* EV36, resulting in fluorescent labelling of whole bacteria, only the signal obtained from ManNAc chemical reporters could be attributed to their incorporation into the polysialic acid capsule. This allowed us to refine our understanding of the capsule metabolic pathways. The method was miniaturized as a microplate assay amenable to screening approaches. With the ability to track the K1 capsule biosynthesis, this platform might be a useful tool for future studies aiming at impacting capsule expression, which is of great interest in the context of increasing pathogen resistance.

## Author Contributions

V.R. and Y.R. performed the experiments. V.R., Y.R. and C.L. analysed and interpreted the data. V.R., Y.R. and C.L. wrote the manuscript. V.R. and C.L. prepared figures. C.L. and C.B. conceptualised the project and acquired funding. C.B., C.L., and Y.R. revised the paper.

## Acknowledgements

This work was supported by the CNRS and the University of Lille. V.R. is a recipient of a research grant from ANR NEURAPROBE (ANR-18-CE07-0042) (C.B.). We thank Dr Corentin Spriet (Plateformes Lilloises en Biologie et Santé (PLBS) - UAR 2014 - US 41) and Dr Boris Vauzeilles for helpful discussions, Pr Antonia P. Sagona for providing us the EV36 strain, and Dr Olivier Vidal for training of V.R.

## SUPPORTING INFORMATION

**Figure S1.**
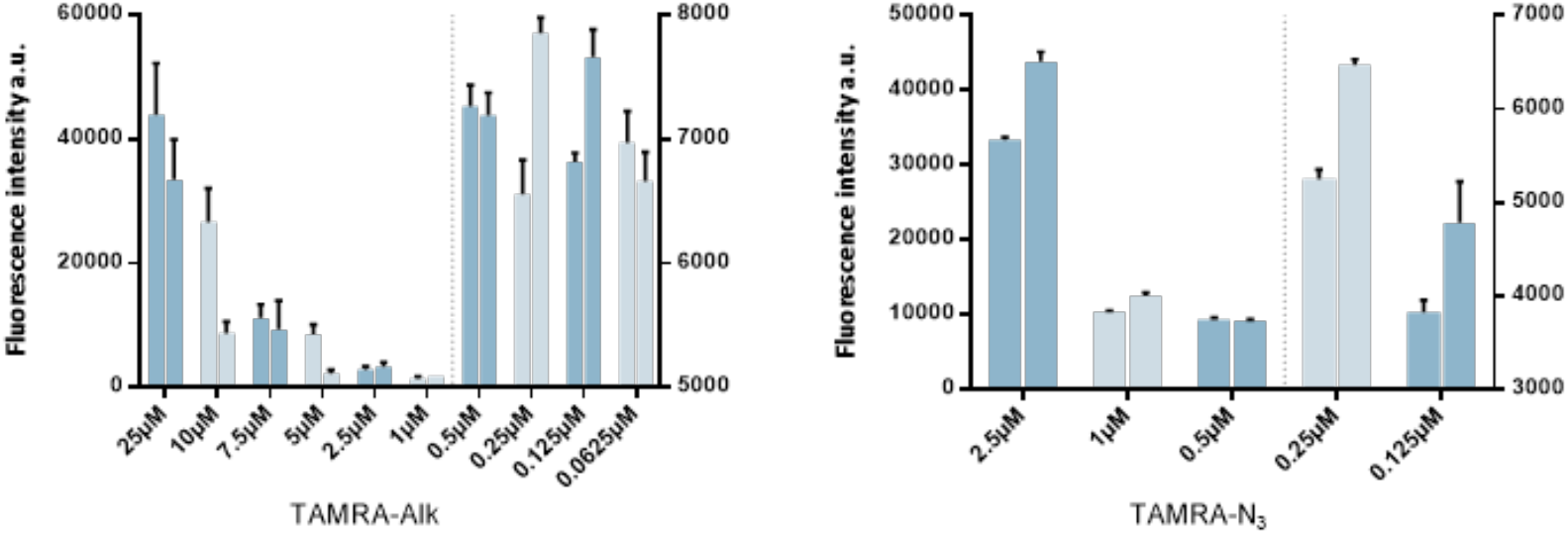
CuAAC optimization.

**Figure S2.**
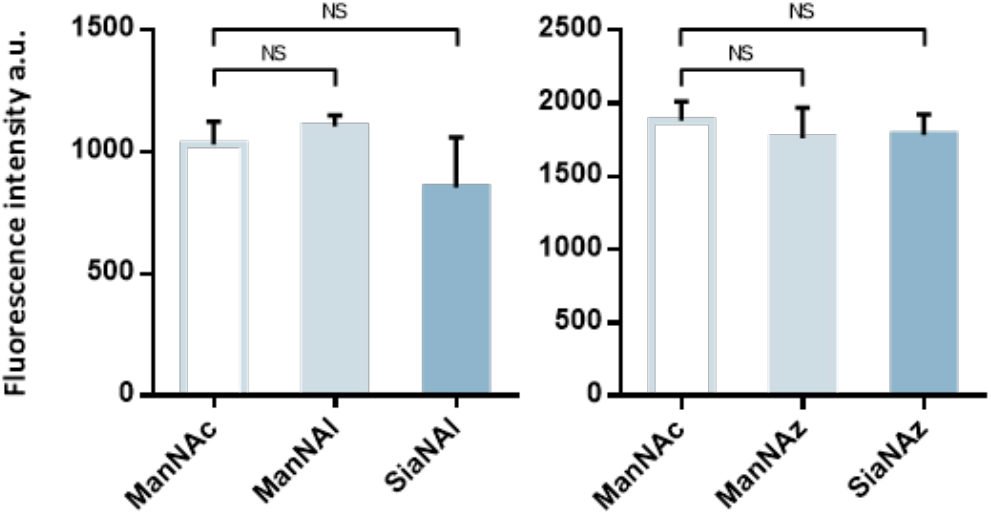
Reporter incorporation in E. coli BL21, a B strain that does not express capsules. Reporters were incubated at 600μM overnight followed by CuAAC with the adequate azide- or alkyne-functionalized TAMRA derivative.

**Figure S3.**
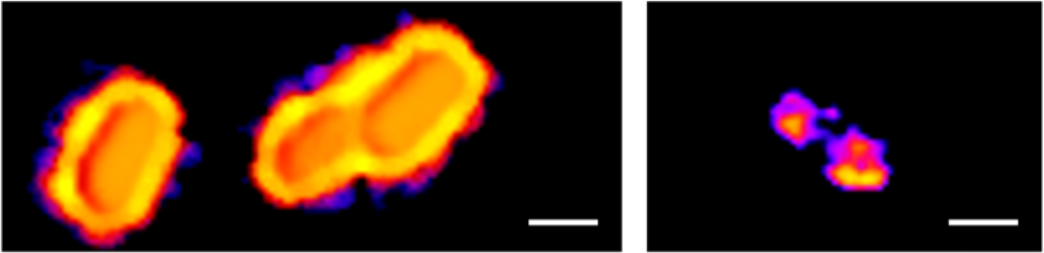
fluorescence microscopy on K-235 bacteria incubated with ManNAz and reacted with TAMRA-Alk. Zooms. **(left)** bacteria exhibiting a peripheral pattern typical of capsule expression, **(right)** bacterium with labelling in the polar regions during cell division.

## References

1 K. S. Kim, Nat. Rev. Neurosci., 2003, 4, 376–385.

2 A. Alkeskas, P. Ogrodzki, M. Saad, N. Masood, N. R. Rhoma, K. Moore, A. Farbos, K. Paszkiewicz and S. Forsythe, BMC Infect. Dis., 2015, 15, 449.

3 J. B. Kaper, J. P. Nataro and H. L. T. Mobley, Nat. Rev. Microbiol., 2004, 2, 123–140.

4 K. J. Kim, J. W. Chung and K. S. Kim, J. Biol. Chem., 2005, 280, 1360–1368.

5 K. S. Kim, H. Itabashi, P. Gemski, J. Sadoff, R. L. Warren and A. S. Cross, J. Clin. Invest., 1992, 90, 897–905.

6 J. B. Robbins, G. H. McCracken, E. C. Gotschlich, F. Ørskov, I. Ørskov and L. A. Hanson, N. Engl. J. Med., 1974, 290, 1216–1220.

7 J. Sarowska, B. Futoma-Koloch, A. Jama-Kmiecik, M. Frej-Madrzak, M. Ksiazczyk, G. Bugla-Ploskonska and I. Choroszy-Krol, Gut Pathog., 2019, 11, 10.

8 B. R. Kunduru, S. A. Nair and T. Rathinavelan, Nucleic Acids Res., 2016, 44, D675–D681.

9 S. M. Steenbergen and E. R. Vimr, Mol. Microbiol., 2008, 68, 1252–1267.

10 B.-X. Lin, Y. Qiao, B. Shi and Y. Tao, Appl. Microbiol. Biotechnol., 2016, 100, 1–8.

11 O. T. Keppler, P. Stehling, M. Herrmann, H. Kayser, D. Grunow, W. Reutter and M. Pawlita, J. Biol. Chem., 1995, 270, 1308–1314.

12 V. V. Rostovtsev, L. G. Green, V. V. Fokin and K. B. Sharpless, Angew. Chemie Int. Ed., 2002, 41, 2596–2599.

13 C. W. Tornøe, C. Christensen and M. Meldal, J. Org. Chem., 2002, 67, 3057–3064.

14 E. M. Sletten and C. R. Bertozzi, Angew. Chemie Int. Ed., 2009, 48, 6974–6998.

15 J. A. Prescher and C. R. Bertozzi, Nat. Chem. Biol., 2005, 1, 13–21.

16 V. Rigolot, C. Biot and C. Lion, Angew. Chemie Int. Ed., 2021, 60, 23084–23105.

17 K. Lang and J. W. Chin, Chem. Rev., 2014, 114, 4764–4806.

18 N. Banahene, H. W. Kavunja and B. M. Swarts, Chem. Rev., 2022, 122, 3336–3413.

19 H. Liang, K. E. DeMeester, C.-W. Hou, M. A. Parent, J. L. Caplan and C. L. Grimes, Nat. Commun., 2017, 8, 15015.

20 A. R. Brown, K. A. Wodzanowski, C. C. Santiago, S. N. Hyland, J. L. Follmar, P. Asare-Okai and C. L. Grimes, ACS Chem. Biol., 2021, 16, 1908–1916.

21 P. Kaewsapsak, O. Esonu and D. H. Dube, ChemBioChem, 2013, 14, 721–726.

22 A. Dumont, A. Malleron, M. Awwad, S. Dukan and B. Vauzeilles, Angew. Chemie Int. Ed., 2012, 51, 3143–3146.

23 J. Mas Pons, A. Dumont, G. Sautejeau, E. Fugier, A. Baron, S. Dukan and B. Vauzeilles, Angew. Chemie Int. Ed., 2014, 53, 1275–1278.

24 N. Geva-Zatorsky, D. Alvarez, J. E. Hudak, N. C. Reading, D. Erturk-Hasdemir, S. Dasgupta, U. H. von Andrian and D. L. Kasper, Nat. Med., 2015, 21, 1091–1100.

25 J. E. Hudak, D. Alvarez, A. Skelly, U. H. von Andrian and D. L. Kasper, Nat. Microbiol., 2017, 2, 17099.

26 P. A. Gilormini, C. Lion, D. Vicogne, T. Levade, S. Potelle, C. Mariller, Y. Guérardel, C. Biot and F. Foulquier, Chem. Commun., 2016, 52, 2318–2321.

27 P. A. Gilormini, C. Lion, D. Vicogne, Y. Guérardel, F. Foulquier and C. Biot, J. Inherit. Metab. Dis., 2018, 41, 515–523.

28 K. N. Chuh, B. W. Zaro, F. Piller, V. Piller and M. R. Pratt, J. Am. Chem. Soc., 2014, 136, 12283–12295.

29 K. Qin, H. Zhang, Z. Zhao and X. Chen, J. Am. Chem. Soc., 2020, 142, 9382–9388.

30 E. R. Vimr and F. A. Troy, J. Bacteriol., 1985, 164, 854–860.

31 C. Møller-Olsen, T. Ross, K. N. Leppard, V. Foisor, C. Smith, D. K. Grammatopoulos and A. P. Sagona, Sci. Rep., 2020, 10, 8903.

32 C. Møller-Olsen, S. F. S. Ho, R. D. Shukla, T. Feher and A. P. Sagona, Sci. Rep., 2018, 8, 17559.

33 J. Plumbridge and E. Vimr, J. Bacteriol., 1999, 181, 47–54.

34 S. K. Behera, A. B. Praharaj, B. Dehury and S. Negi, Glycoconj. J., 2015, 32, 575–613.

35 Y. Talyansky, T. B. Nielsen, J. Yan, U. Carlino-Macdonald, G. Di Venanzio, S. Chakravorty, A. Ulhaq, M. F. Feldman, T. A. Russo, E. Vinogradov, B. Luna, M. S. Wright, M. D. Adams and B. Spellberg, PLOS Pathog., 2021, 17, e1009291.

36 S. S. Spicer and L. Warren, J. Histochem. Cytochem., 1960, 8, 135–137.

37 B. Revilla-Nuin, A. Reglero, M. A. Ferrero and L. B. Rodrıguez-Aparicio, FEBS Lett., 1999, 449, 183–186.

38 E. R. Vimr, ISRN Microbiol., 2013, 2013, 1–26.

39 M. Petersen, W.-D. Fessner, M. Frosch and E. LÃ¼neberg, FEMS Microbiol. Lett., 2000, 184, 161–164.

40 C. Brigham, R. Caughlan, R. Gallegos, M. B. Dallas, V. G. Godoy and M. H. Malamy, J. Bacteriol., 2009, 191, 3629–3638.

41 E. R. Vimr, K. A. Kalivoda, E. L. Deszo and S. M. Steenbergen, Microbiol. Mol. Biol. Rev., 2004, 68, 132–153.

42 M. Sawa, T.-L. Hsu, T. Itoh, M. Sugiyama, S. R. Hanson, P. K. Vogt and C.-H. Wong, Proc. Natl. Acad. Sci., 2006, 103, 12371–12376.

43 C. Whitfield, S. S. Wear and C. Sande, Annu. Rev. Microbiol., 2020, 74, 521–543.

44 M. X. Henriques, T. Rodrigues, M. Carido, L. Ferreira and S. R. Filipe, Mol. Microbiol., 2011, 82, 515–534.

45 S. Phanphak, P. Georgiades, R. Li, J. King, I. S. Roberts and T. A. Waigh, Langmuir, 2019, 35, 5635–5646.

46 F. W. Studier and B. A. Moffatt, J. Mol. Biol., 1986, 189, 113–130.

47 S. Pelkonen, J. Häyrinen and J. Finne, J. Bacteriol., 1988, 170, 2646–2653.

