## Supplementary figures and images for "A bioorthogonal chemistry approach to detect the K1 polysialic acid capsule in *Escherichia coli*"

### Figure S1

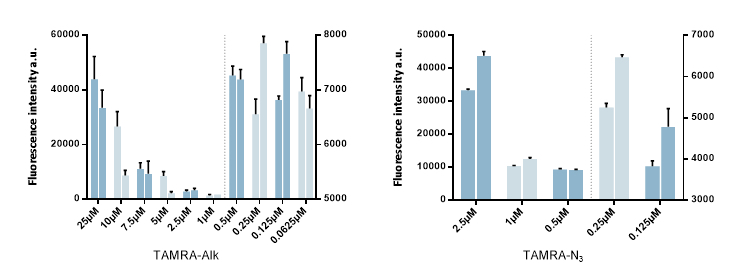

### Figure S2

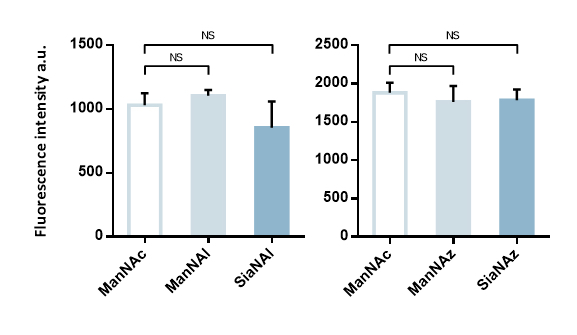

### Figure S3

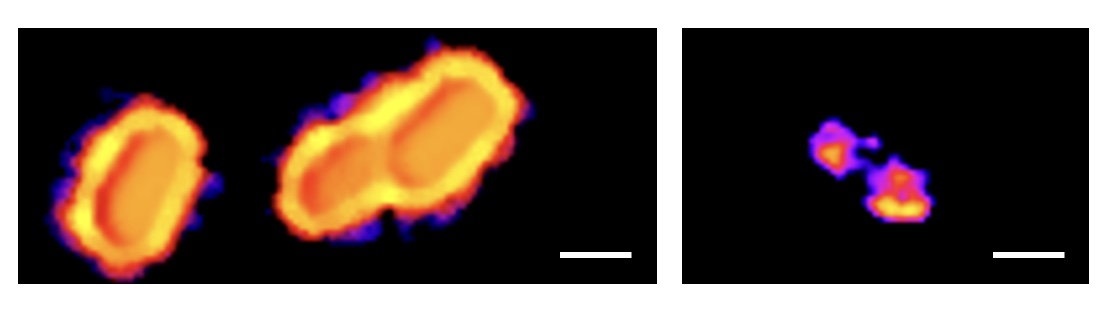
